# Population-level transposable element expression dynamics influence trait evolution in a fungal crop pathogen

**DOI:** 10.1101/2023.03.29.534750

**Authors:** Leen Nanchira Abraham, Ursula Oggenfuss, Daniel Croll

**Author notes:** Present address: Institute of Plant Sciences, University of Cologne, Cologne, Germany (Leen Nanchira Abraham). Present address: Department of Microbiology and Immunology, University of Minnesota Medical School, Minneapolis, Minnesota, United States of America (Ursula Oggenfuss).

## Abstract

Rapid adaptive evolution is driven by strong selection pressure acting on standing genetic variation within populations. How adaptive genetic variation is generated within species and how such variation influences phenotypic trait expression is often not well understood though. Here, we focused on recent activity of transposable elements (TEs) using deep population genomics and transcriptomics analyses of a fungal plant pathogen with a highly active content of TEs in the genome. *Zymoseptoria tritici* causes one of the most damaging diseases on wheat, with recent adaptation to the host and environment being facilitated by TE-associated mutations. We obtained genomic and RNA-sequencing data from 146 isolates collected from a single wheat field. We established a genome-wide map of TE insertion polymorphisms in the population by analyzing recent TE insertions among individuals. We quantified the locus-specific transcription of individual TEs within the population and revealed considerable variation in transcription across individual TE loci. About 20% of all TE copies show activity in the genome implying that genomic defenses such as repressive epigenetic marks and repeat-induced polymorphisms are ineffective at preventing the proliferation of TEs in the genome. A quarter of recent TE insertions are associated with expression variation of neighboring genes providing broad potential to influence trait expression. We indeed found that TE insertions are likely responsible for variation in virulence on the host and secondary metabolite production. Our study emphasizes how TE-derived polymorphisms segregate even in individual populations and broadly underpin transcription and adaptive trait variation in a species.

## Introduction

Rapid adaptive evolution enables species to cope with challenging environmental conditions including climate change. Most evidence for rapid evolutionary processes comes from experimental studies applying artificial selection pressures (Barrett & Schluter, 2008). In natural populations, the extent of segregating adaptive genetic variation is central for the evolutionary response to environmental change. Transposable elements (TEs) were recently recognized as key drivers of genetic variation within species and even single populations (Cai et al., 2022a; De Kort et al., 2022; Oggenfuss et al., 2021). TEs are genomic sequences that can be mobilized, and high activity in some species was linked to the gain of adaptive variation (Britten, 2010; Chuong et al., 2017; Hosaka & Kakutani, 2018; Wei et al., 2022a). Based on the mode of proliferation, TEs are categorized into retrotransposons, which copy via an RNA intermediate, and DNA transposons which can excise and integrate into a different locus. TE-encoded transposase proteins mediate the excision and integration of TEs into the genome (Suh, 2019). TE transposition can lead to insertional mutations that disrupt *cis*-regulatory elements or serve as alternative promoters that render the gene more responsive (Sundaram & Wysocka, 2020; Ullastres et al., 2021). Similarly, TE exaptation into the coding sequence of a genome can produce novel regulatory sequences and lead to intronization or exonization events with beneficial or detrimental effects for the host (Cosby et al., 2020; Etchegaray et al., 2021). Also, TEs can be an important source of epigenetic modifications regulating gene expression (Young et al., 2020). Thus, TE-induced genetic variation has vast potential to alter the genomic architecture and modulate phenotypic trait expression.

Genetic variation within species is generated to a significant extent by TEs actively creating new copies (Almeida et al., 2022; Feschotte, 2008; Slotkin & Martienssen, 2007a). Active TEs can also cause instabilities in chromatin and genomic rearrangements (Baduel & Quadrana, 2021). However, most TE copies in eukaryotic genomes are not transcriptionally active, despite making up a large fraction of the genome (Slotkin & Martienssen, 2007b). TE accumulation in genomes may be harmful, favoring the evolution of defense mechanisms. TE-rich genomic regions carry repressive epigenetic modifications, such as histone modifications and DNA methylation. These modifications prevent the transcriptional machinery from accessing TE sequences, thereby silencing TE copies. A prominent defense mechanism against active transposition is RNA interference, which can bind and cleave transcribed TE sequences (Bucher et al., 2012; Mita & Boeke, 2016). Beyond RNA-based defenses, repeat-induced point mutations (RIP) are a silencing mechanism specific to fungi that induces C->T mutations in repetitive DNA. All defense mechanisms are expected to be highly conserved to prevent the uncontrolled proliferation of TEs. Assessing the transcriptional landscape of TEs in the genome helps distinguish between silenced and deactivated TEs from copies with the ability to transpose. Assessing the expression state of individual copies of the same TEs is challenging when using short read sequencing datasets, because the mapping to individual copies is often ambiguous (Berrens et al., 2021; Lanciano & Cristofari, 2020). Evolutionarily young TEs with high recent activity are particularly challenging, as the genome may contain many near identical copies. Advances in sequencing technologies and mapping approaches have overcome some limitations to identify locus-specific expression profiles of different TEs and helped uncover new mechanisms of TE regulation (Berrens et al., 2021; Schwarz et al., 2022; Yang et al., 2019).

Active TEs generate TE insertion polymorphisms (TIPs) within species with individual genotypes differing in their presence or absence of TEs across loci. Genome-wide association studies in tomatoes identified 40 TIPs robustly associated with phenotypic trait variation (Domínguez et al., 2020). In accessions of *Arabidopsis thaliana,* thousands of recent transposition events were discovered with the potential to facilitate climatic and other adaptations (Quadrana et al., 2016). Similarly, 1,059 *Oryza sativa* genomes revealed an association of miniature inverted-repeat transposable elements (MITEs) with agronomically important traits such as grain width (Carpentier et al., 2019). TIPs can also contribute to changes in gene expression by attracting epigenetic marks to neighboring genes (Chen et al., 2020). Genetic variation originating from TE insertion dynamics is an abundant source of genomic variation in plant pathogenic fungi. The distribution of TEs in fungal genomes can alter gene expression including the expression of small secreted proteins acting as virulence factors, also called effectors. The soil-borne fungal plant pathogen, *Verticillium dahliae* carries a dynamic TE-rich region encoding virulence-related genes (Ramírez-Tejero et al., 2020; Torres et al., 2021). TEs also played a key role in genome plasticity and induced chromosomal rearrangement in the rice blast pathogen *Magnaporthe oryzae*. A strong association of TEs and secreted proteins involved in virulence was observed in a genomic region enriched in repeats and virulence genes (Bao et al., 2017; Jeon et al., 2015; Zhang et al., 2021). Effector genes in the plant pathogen *Leptosphaeria maculans* were found in TE-rich heterochromatic regions (Soyer et al., 2014). TE insertions modulated the epigenetic state of nearby effector genes, thereby modulating their expression patterns. *Pyrenophora tritici-repentis,* a necrotrophic fungal pathogen of wheat, showed higher TE content in pathogenic isolates compared to nonpathogenic isolates, with evidence of TE-mediated effector diversification and movement of virulence factors (Gourlie et al., 2022; McDonald et al., 2019; Moolhuijzen et al., 2018). Existing evidence highlights the vast potential of TE-mediated genetic, epigenetic, and phenotypic changes occurring in rapidly evolving pathogen populations.

*Zymoseptoria tritici* is an economically important fungal wheat pathogen with a genome harboring 16-24% TEs (Badet et al., 2020). The pathogen has a repertoire of virulence factors to manipulate the host plant (Kilaru et al., 2022; Mirzadi Gohari et al., 2015; Morais do Amaral et al., 2012). Effector gene expression can change the epigenetic landscape of TEs during plant infection (Fouché et al., 2020). Silenced TEs near an important effector are linked to reduced effector expression leading to more damage on a specific wheat cultivar (Fouché et al., 2020; Meile et al., 2018, 2020). TE insertions upstream of the gene encoding the transcription factor Zmr1 regulate melanin production, a pigment required for pathogen survival during stressful conditions (Krishnan et al., 2018). The insertion of a retrotransposon into the promoter of a gene encoding a transporter increases fungicide efflux and contributes to multi-drug resistance (Omrane et al., 2017). Similarly, TE insertions are linked to the adaptive deletion of a gene encoding an effector, which is likely recognized by the plant host (Hartmann et al., 2017). The activity of TEs is generally high in the pathogen and varies with geography (Badet et al., 2020; Feurtey et al., 2023; Oggenfuss et al., 2021; Oggenfuss & Croll 2023).

In this study, we interrogated large transcriptomic, genomic, and phenotypic datasets of a *Z. tritici* population to identify associations between active TEs and adaptive trait variation. We further analyzed variation in TE activity among individuals and quantified the strength of genomic defenses against TE loci and repressive epigenetic marks. Finally, to capture TE- driven phenotypic variation in the pathogen, we associated metabolite production variation and pathogenicity-related traits of the fungal population with TE insertion polymorphisms.

## Materials and methods

### Fungal isolates and sequencing

We analyzed *Z. tritici* isolates collected from a wheat field in Eschikon, Switzerland. Wheat cultivars were planted in a randomized block design and infections occurred naturally from local inoculum or adjacent fields (Singh et al., 2021). Total genomic DNA was extracted from 139 *Z. tritici* isolates yeast-sucrose broth (YSB) cultures using the QIAGEN DNAeasy Plant Mini Kit. Illumina libraries were prepared using the TruSeq Nano DNA Library Prep kit. Sequencing was performed in 100 bp paired-end mode on a HiSeq 4000 at the iGE3 sequencing platform (Geneva, Switzerland). Raw reads are available on the NCBI Short Read Archive under the BioProject PRJNA596434. For RNA sequencing, the same isolates (146 isolates, were cultured in a Vogel’s Medium N (Minimal) modified as ammonium nitrate replaced with potassium nitrate and ammonium phosphate (Perkins, 2006) without sucrose and agarose to induce hyphal growth. Total RNA was isolated from the filtered mycelium after 10-15 days using the NucleoSpin® RNA Plant and Fungi kit. The RNA concentrations and integrity were checked using a Qubit 2.0 Fluorometer and an Agilent 4200 TapeStation system, respectively. Only high-quality RNA (RIN>8) was used to prepare TruSeq-stranded mRNA libraries with a 150 bp insert size. Sequencing was performed on an Illumina HiSeq 4000 in 100 bp single-end mode. Raw reads for RNAseq are available on the NCBI Short Read Archive under the BioProject PRJNA650267 (Abraham & Croll 2022; Supplementary Table S1).

### Identification of TE insertion polymorphism in Z. tritici

Whole-genome Illumina sequencing reads were quality checked using FastQC (version 0.11.5, (*FastQC: A Quality Control Tool for High Throughput Sequence Data – ScienceOpen, n.d.*)) and trimmed with Trimmomatic version 0.36 (Bolger et al., 2014) to remove adapter sequences and low-quality reads with parameters ILLUMINACLIP: TruSeq3-PE.fa:2:30:10 LEADING:3 TRAILING:3 SLIDING WINDOW:4:15 MINLEN:36. To detect TE insertions, we used the R based tool *ngs_te_mapper* version 79ef861f1d52cdd08eb2d51f145223fad0b2363c (Linheiro & Bergman, 2012) integrated into the McClintock pipeline version 20cb912497394fabddcdaa175402adacf5130bd1 (Nelson et al., 2017). Filtering and validation of the identified insertion polymorphisms followed previously established protocols for the same species (Oggenfuss et al., 2021). To confirm the presence of predicted non-reference TEs, we extracted the reads mapped near the predicted insertion site and assessed whether the target site duplication represented a break in the alignment as expected. Using spliced junction reads, we analyzed if a gap was suggested in the region of the position indicating the absence of a reference genome TE copy in a particular isolate.

### Locus-specific TE and gene expression analyses

Locus-specific expression profiles for TEs were generated using mapped RNA-seq data with the tool SQuIRE (v0.9.9.9a-beta) (Yang et al., 2019). TE annotation of the reference genome IPO323 was retrieved from (Badet et al., 2020). The SQuIRE “Map” mode was used to align RNA-seq data and SQuIRE “Count” with the –EM parameter to perform the estimation-maximization algorithm to quantify TE expression considering both uniquely mapped and multi-mapped reads typically found in repetitive sequences such as TEs. TE-derived reads normalized by fragments per kilobase of exon per million (FPKM) were filtered to keep the reads originating from annotated strand direction in the IPO323 reference genome. In parallel, gene expression profiles were analyzed using QTLtools (version 1.1, (Delaneau et al., 2017)). RNA-seq reads mapped to the IPO323 reference genome were analyzed based on high-quality gene models (Grandaubert et al. 2015). Read counts were summarized with QTLtools -- quan mode. Only reads with a minimum Phred mapping quality > 10 were kept for further analyses. Reads were normalized using the --rpkm (reads per kilobase of transcript per million reads mapped) from QTLtools (version 1.1) (Delaneau et al., 2017).

### Association mapping analyses for expression variation

SNPs used for the association mapping were retrieved from a variant calling procedure performed with raw sequencing reads checked by FastQC (version 0.11.5) (Andrews, 2010) and trimmed with Trimmomatic Click or tap here to enter text.to remove adapter sequences and low-quality reads with parameters ILLUMINACLIP: TruSeq3-PE.fa:2:30:10 LEADING:3 TRAILING:3 SLIDING WINDOW:4:15 MINLEN:36. Trimmed sequences were aligned to the *Z. tritici* IPO323 reference genome (see above) using bowtie2 version 2.3.4.3 (Langmead & Salzberg, 2012) with the option --very-sensitive-local. Variant calling was performed using Haplotypecaller integrated in the Genome Analysis Toolkit (GATK) v. 4.0.11.0 (McKenna et al., 2010). We retained SNPs with QUAL>1000, AN>20, QD>5.0, MQ>20.0, ReadPosRankSum_lower=2.0, ReadPosRankSum_upper=2.0, MQRankSum_lower=2.0, MQRankSum_upper=2.0, BaseQRankSum_lower=2.0, BaseQRankSum_upper=2. Variants passing quality filtration were further filtered to remove multiallelic SNPs using the *bcftools* (version 1.9) --norm option. Variants were filtered to keep only variants genotyped in at least 90% of the individuals and common variants ≥5% using the *VCFtools* --max-missing and *bcftools* -q 0.05 minor option (Danecek et al., 2011a, 2021). Expression quantitative trait loci (eQTL) for variation in locus-specific TE expression were searched using QTLtools (version 1.1) (Delaneau et al., 2017) in the *cis* --permutation mode with 1000 permutations and 5 kb *cis* windows surrounding each TE locus. The permutation *p*-values were false discovery rate (FDR) corrected to identify the top eQTL (5% threshold). We also analyzed gene expression variation associated with TIPs. For this, only genes within 5 kb of each TIP were tested for associations with gene expression levels. Fold change in gene expression was calculated from the ratio of mean gene expression between isolates with TE insertion and isolates without TE insertion. A Wilcoxon test was performed to assess significance of the association.

### Phenotype-genotype association mapping for TIPs

We analyzed genome-wide TIPs for associations with phenotypic trait variation (*i.e.*, TIP-GWAS). To assess variation in virulence phenotype variation, we performed an infection assay on the Swiss winter wheat cultivar Claro grown in a growth chamber (Singh et al., 2021). We used diluted (2 × 10^5^ spores/ml in 15 ml of sterile water containing 0.1% TWEEN2) 8-day old YSB-grown *Z. tritici* spore suspension to infect the three weeks old wheat plant. After spray inoculation, the plants were kept at 100% humidity for 21 days. Leaf lesions were assessed using ImageJ. the ratio of the total lesion area and total leaf area was calculated to obtain the percent leaf area covered by lesions (PLACL) (Karisto et al., 2018).

The metabolome composition of the pathogen population was assayed previously using untargeted metabolite profiling based on UPLC-HRMS (Singh et al., 2022). *Z. tritici* isolates grown in YSB for 8 days were filtered through cheesecloth to remove hyphae and washed in milli-Q water to remove media traces. The spores were suspended and lyophilized to extract metabolites in 1 ml of HPLC-grade methanol. The extract was centrifuged at 15,000 rpm for 5 min to pellet down debris and this last step was repeated until a clear supernatant was recovered. Untargeted metabolite profiling was carried out by UHPLC-HRMS using an Acquity UPLC coupled to a Synapt G2 QTOF mass spectrometer (Waters, Inc.). Formic acid (0.05%) in water as mobile phase A and formic acid (0.05%) in acetonitrile as mobile phase B with a gradient of 0–100 % B in 10 min, holding at 100% B for 2.0 min, re-equilibration at 0% B for 3.0 min was used. Samples were analyzed using mass spectrometric parameters of 50- 1200 Da, 0.2 s scan time, 120°C source temperature, 2.5 kV capillary voltage, 25V cone voltage, 900 L/h desolvation gas flow, and 400°C, 20 L/h cone gas flow, and 4 eV collision energy (low energy acquisition function) or 15–50 eV collision energy (high energy acquisition function). Recordings were made using Masslynx XS v.4.1 (Waters Inc.). Detecting markers with Markerlynx XS was performed with the following parameters: initial and final retention times 1.5 and 10 minutes, mass range 85–1200 Da, mass window 0.02 Da, retention time window 0.08 min, intensity threshold 500 counts, automatic peak width calculations, de-isotoping applied. We used untransformed relative abundance values for each peak from the metabolome analysis for metabolome variation association mapping.

TE insertion loci (TIPs) with a minor allele frequency of >5% (for either the presence or absence of the TE at the locus) were used for association mapping with phenotypic trait variation based on mixed linear models and performs likelihood ratio tests (--lmm 2) with the GEMMA version 0.98.3) (Zho & Stephens, 2012). The standardized relatedness matrix calculated from individual genotypes was used to correct for uneven relatedness in association mapping. Association *p-*values were considered significant using Bonferroni (Bland & Altman, 1995) multiple comparison corrections. The Bonferroni threshold was calculated by dividing the nominal threshold of ⍺ = 0.05 by the total number of TIPs used in our GWAS (*n* = 192). Additionally, we considered a 5% false discovery rate (FDR) threshold using the *p.adjust* function in the R package *stat* (Korthauer et al., 2019). Linkage disequilibrium (*r*^2^) between TIPs and SNPs in a 5kb upstream and downstream distance from the TIP locus were calculated using the “–hap-r2” in VCFtools v. 0.1.15 (Danecek et al., 2011b). Linkage disequilibrium analyses between TIP alleles (presence/absence) and calculated the *r^2^* for all the SNPs in the window to the TIP locus polymorphism. We also calculated *r*^2^ for all pairwise SNP combinations in the same windows.

## Results

### A diverse pool of active TEs in the genome of Z. tritici

The genome of the fungal wheat pathogen *Z. tritici* contains 16–24% TEs based on reference-quality genome assemblies (Badet et al., 2020; Goodwin et al., 2011). We assessed the strength of selective constraints against TEs by analyzing the distribution of TE families relative to gene elements (Supplementary Table S2). Based on the telomere-to-telomere assembled reference genome IPO323, we found that ∼10% of TEs in the genome were located within 1 kb upstream of coding regions likely overlapping with regulatory elements (Fig 1A). Similarly, exons (6%), 3’ untranslated regions (UTRs, 6%), and downstream regions (<1 kb, 10%) also carried high proportions of TEs near genes. Introns and 5’ UTRs showed low TE counts (1% and 2%, Fig 1A). Across gene elements, only few DNA transposons inserted in exons, while retrotransposons made up the highest proportion (Fig. 1B). To investigate the distribution of major TE families across gene elements, we focused on the most abundant TE families (≥25 copies in the genome) including five DNA transposons, three retrotransposons and two unassigned TE families (Fig. 1C). We found that 56% of the DNA transposon DTH *Donna* copies were integrated primarily into introns consistent with previous findings of intron sequences being at least partially occupied by TEs in *Z. tritici* (Torriani et al., 2011). Similarly, more than half of the copies of the Tc1-*Mariner* (DTT) superfamily and a *Copia* element (RLC Deimos) were integrated into upstream regions of genes (Fig. 1C). TE families *Gliese*, the MITEs *Troll,* and *Goblin* showed the highest insertion proportions in 3’ UTRs. We next analyzed associations of TE insertions and gene expression levels under nutrient-limited conditions. TEs integrated most frequently close to the 10% lowest expressed genes and integration events were dominated by retrotransposons (Fig. 1D). *Gypsy* (recently renamed as *Ty3*) retrotransposons and unassigned TEs show the strongest skew towards integration close to lowest expressed genes (Supplementary Fig S1). On the contrary, DNA transposons are less depleted near strongly expressed genes. Overall, the TE distribution in the genome is strongly shaped by negative selection and correlates with the transcriptional landscape of genes.

**Figure 1.**
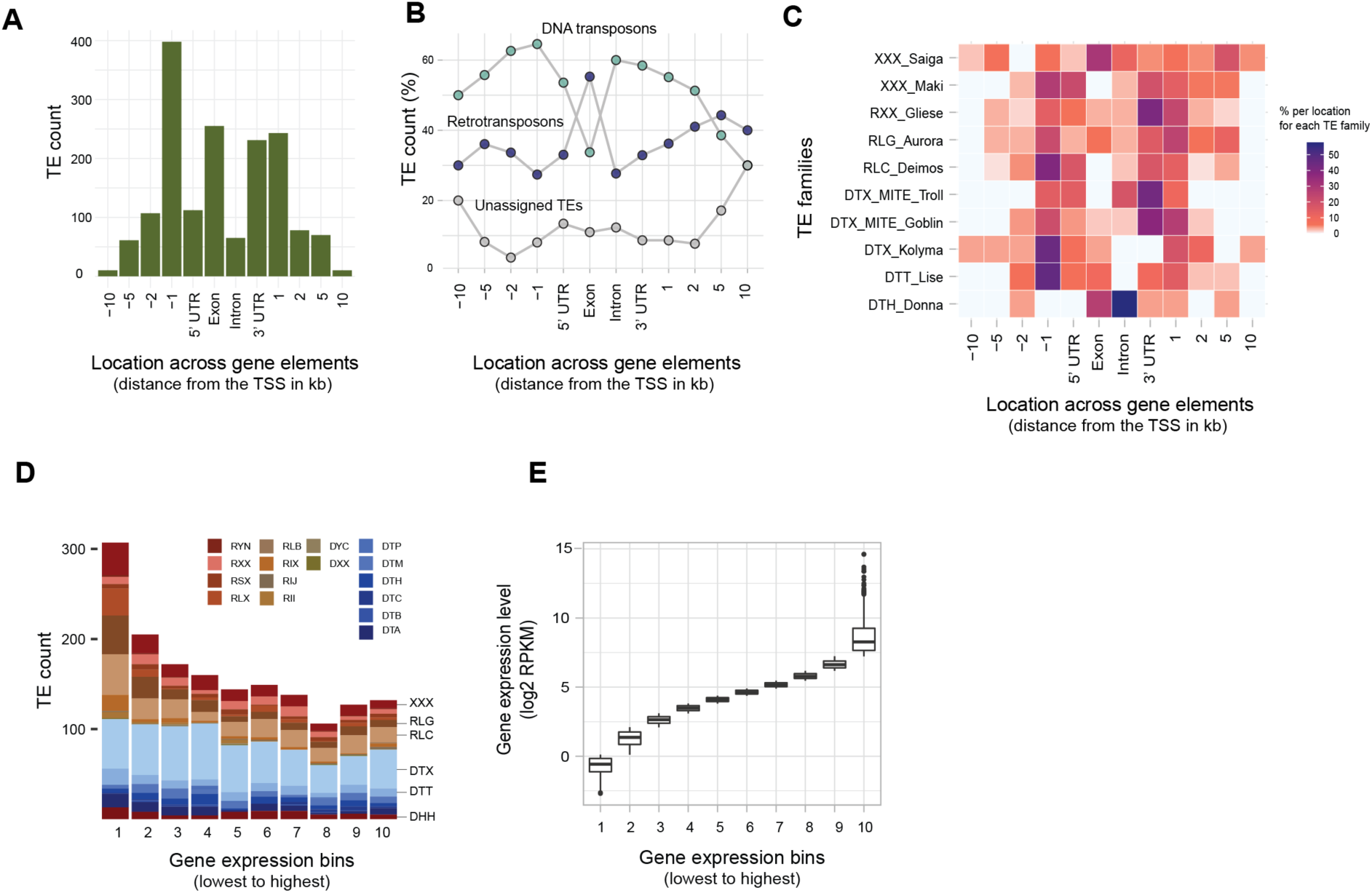
Distribution of transposable elements (TEs) in gene elements in the reference genome of *Zymoseptoria tritici*. (A) Genomic localization of TEs in gene element and 10kb windows upstream and downstream of the transcription start site (TSS). (B) Distribution of DNA and retrotransposon in gene elements and 10kb windows upstream and downstream of TSS. (C) Distribution of different TE families in gene elements and 10kb windows upstream and downstream of TSS. (D) The number of TE insertions close to genes. Gene expression bins refer to genes ordered by increasing order of gene expression and the bins group equal numbers of genes.

### TE insertion dynamics in a field population

TEs in the species show recent activity and contribute to genome-wide polymorphism (Oggenfuss & Croll, 2023). We examined TE generated polymorphism based on TIPs for a deeply sampled single field population of genetically diverse isolates (Supplementary Table S3). We used *ngs_te_mapper* to scan 139 whole-genome sequencing read sets individually aligned to the reference genome. TEs absent in the reference genome were rare in the population (“non-reference TEs”; on average carried by 5% of all isolates) in contrast to TEs with a copy present in the reference genome (“reference TEs”; Fig. 2A). This is consistent with the expectation that TEs present in the reference genome are more likely to be at a higher frequency among other isolates as well. TIR elements (DTX) are the most frequent TE family showing TIPs in the population followed by SINE (RSX) and mutator elements (DTM; Fig. 2C). TIPs inserted into introns had the highest frequency in the population compared to TEs inserted in other gene elements (Fig. 2B). For example, 46% of isolates contained intron insertions of the TIR DTX element. Furthermore, 79% of isolates from the population carried LTR elements (RLX) in exons (Fig. 2D). Given the high numbers of TIPs nearby coding sequences, we analyzed associations of TIPs with gene expression (Supplementary Table S4). We considered genes having TIPs within 10 kb and compared transcript levels between isolates with and without TE insertions. Approximately 25% of the TIPs in the population showed a significant association with the expression of a neighboring gene (*p-*value < 0.05). In general, TE insertions nearby a gene were significantly associated with lower gene transcription levels (Fig. 2E). We further analyzed the population frequency of the inserted TEs at TIPs and the association with gene expression variation among isolates. We found that low-frequency TEs were significantly associated with higher expression variation of neighboring genes (Fig. 2F). Overall, the genomic landscape of recent TE insertions reflects gene expression profiles.

**Figure 2.**
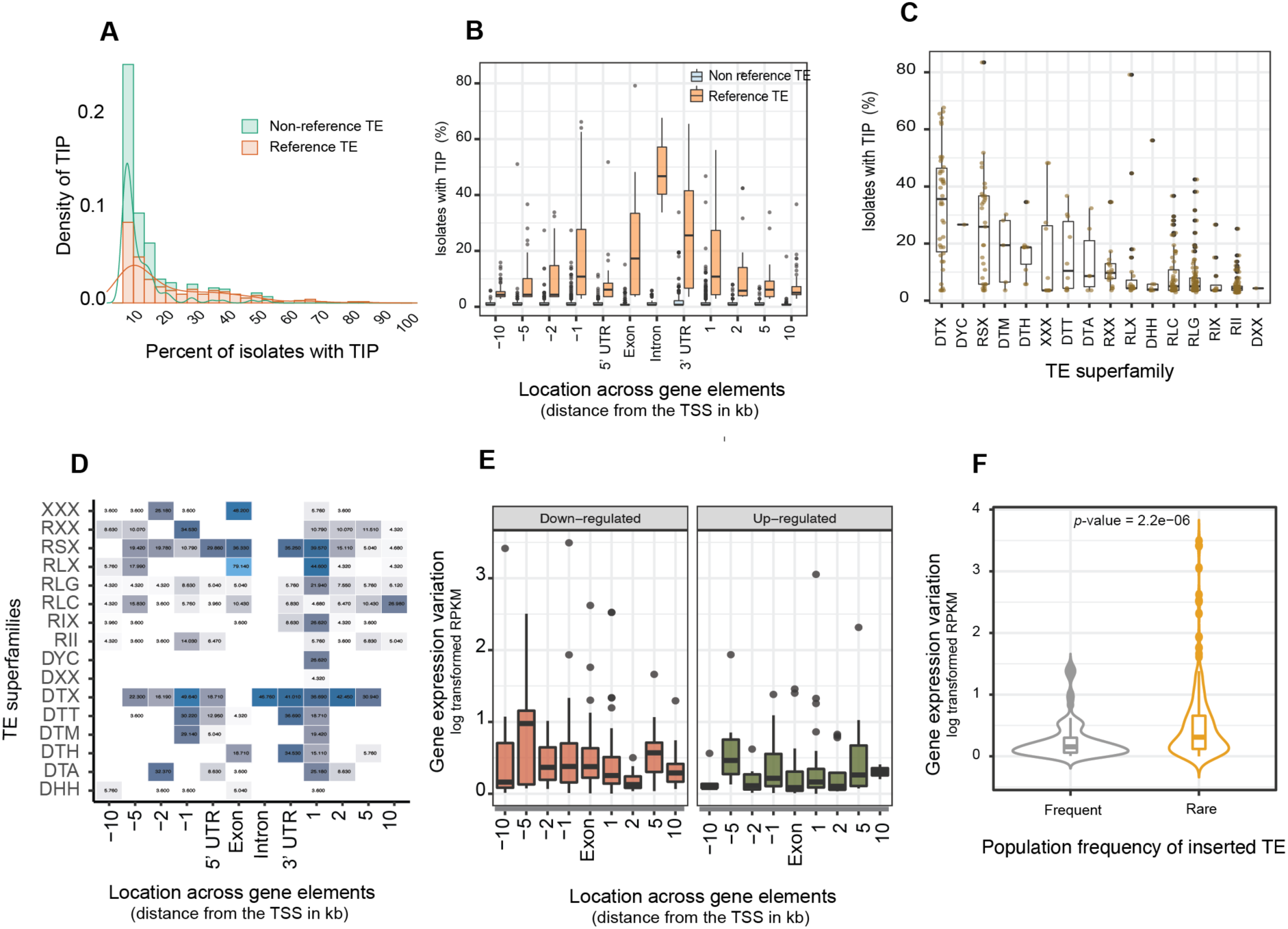
TE insertion polymorphisms in a single field population and association with gene expression variation. (A) Frequency of TEs present in the reference genome (“reference TEs”) and (“non-reference TEs”) (B) Frequency of TE insertion polymorphisms (TIPs) present in different gene elements (C) Frequency of different TE superfamilies in the population D) Frequency of different TE superfamilies in the population according to the gene elements. (E) Association of TIPs with the expression variation of neighboring genes. (F) Gene expression variation for TIPs with high insertion TE frequency (>20 in the population) *vs.* for TIPs with rare TEs (frequency <20).

### Locus-specific expression of TEs

Transcription of TEs can lead to the transposition and copy-number increases. To identify the pool of potentially active TEs from each TE family, we analyzed evidence for transcription across all members of TE families (Supplementary Table S5). We generated locus-specific expression estimates based on uniquely mapping RNA-sequencing reads. Reads mapping to multiple loci in the genome were distributed proportionally across loci according to the algorithm. We found that 43% of all TE copies per genome show evidence of transcription (Supplementary Fig. S2). Sequence similarity of recently transposed TE copies can be a challenge for the unique mapping of sequencing reads to individual loci. Hence, locus-specific analyses are likely biased against the youngest TEs in the genome. To assess this risk, we reconstructed the phylogenetic tree of a MITE family using DNA sequences of individual copies in the genome. We assessed transcript abundance for each locus in association with the terminal branch length of the individual copy as an indicator of age. We found no meaningful association between the age of a TE copy and the estimated transcript abundance suggesting that expression quantification is accurate even for the youngest TE copies (Fig 3A). We observed the highest transcription levels for *Copia* and *Gypsy/Ty3* retrotransposons (Fig 3B,C). Furthermore, we found high variation among individuals in the population for the expression of some DNA transposon TIRs (Fig 3D,E). The MITE family Undine showed high expression variation among isolates followed by the DTA_hAT superfamily (standardized residuals = 8.0 and 7.1, respectively). Although the number of TE copies significantly differs among chromosomes, we found no significant variation in transcription (Supplementary Fig. S3). We found though a higher proportion of TE copies residing within 1 kb upstream of the transcription start sites. We also found that TE copies further away from the transcription start site tended to show comparatively high variation in transcription among individuals compared to TE copies closer to genes.

**Figure 3.**
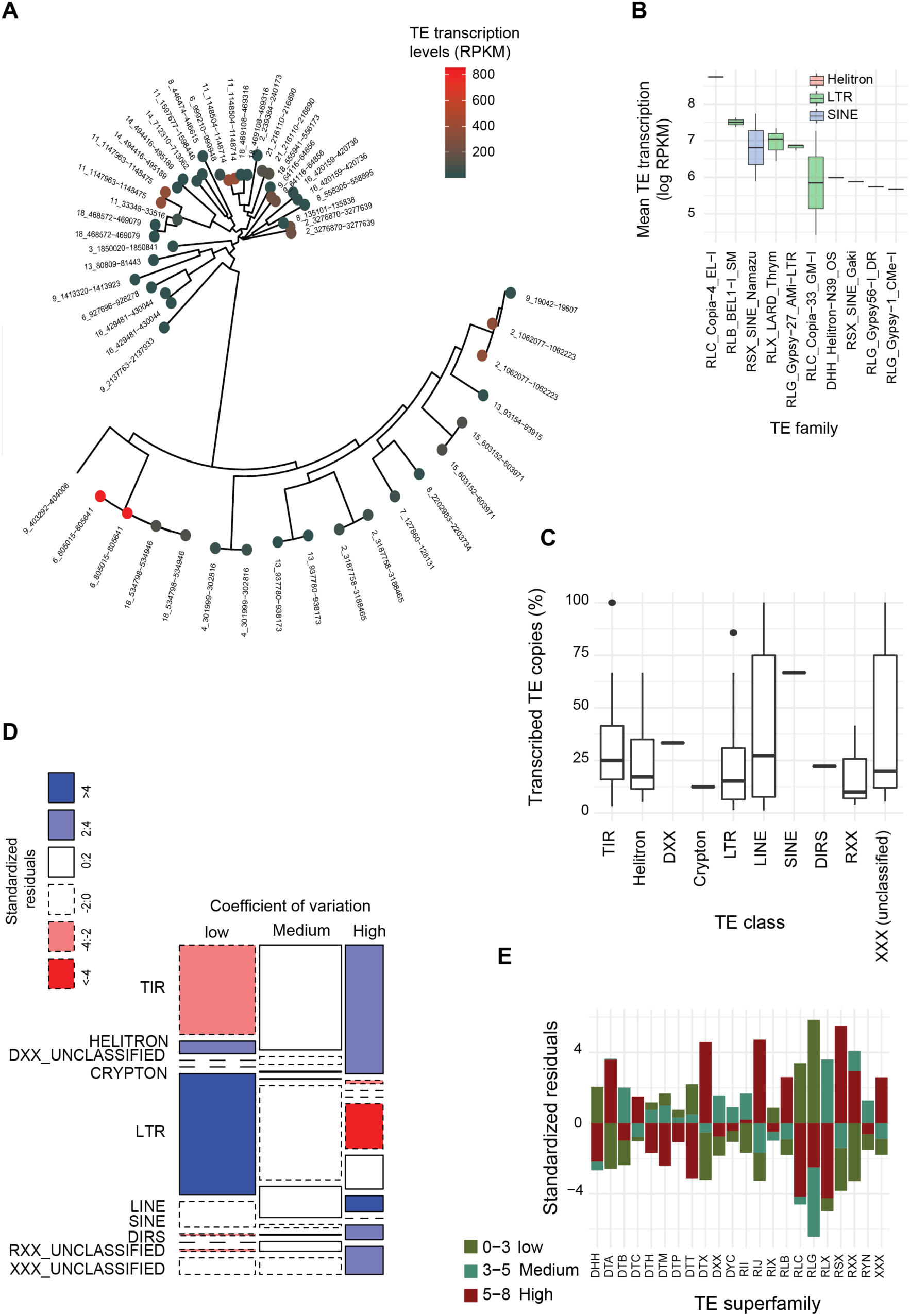
Population-level variation of locus-specific transcription of TEs. A) Phylogenetic tree of all copies of a MITE family discovered in the IPO323 reference genome. Colors highlight transcription levels for each copy. B) Transcription levels of individual copies of different TE families in the genome (ten most transcribed TE families). C) Percent of TE loci showing transcription in the population. D) Likelihood of different TE classes with different categories of transcriptional variation (0-3: low coefficient of variation; 3-5: medium coefficient of variation, 5-8: high coefficient of variation). E) Standardized residuals of coefficient of variation categories.

The genomic context as well as the identity of individual TEs are significant factors explaining genome-wide TE transcription and insertion activity (Fig 4A). Epigenetic factors such as histone methylation marks including H3K4me2, H3K9me2, H3K9me3, and H3K27me3 can have important effects on gene and TE transcription. In addition, repeat-induced point (RIP) mutations, and GC content of TE loci and their association with transcription can be important indicators of the activity of TEs. We found a negative correlation between TE transcription levels and counts of RIP mutations. This is consistent with the action of RIP as a genomic defense against TEs. GC content, repressive histone methylation marks (H3K27m3 and H3K9m2), and the age of the TE were positively correlated with TE transcription (Fig. 4B). Surprisingly, also the euchromatic H3K4m3 mark was positively correlated with higher TE transcription. The non-autonomous TE family DTX MITE Undine showed a negative correlation between transcription and the age of the TE copy (using terminal branch lengths) and a positive correlation with epigenetic markers such as repressive and constitutive heterochromatin. Transcription of the DNA transposon DTA hAT copies showed a positive correlation with repressive H3K9me2 marks (Fig. 4C). We further investigated interactions of the genomic environment with TE activity. We found a positive relationship between the proportion of transcribed copies of a TE in a genome with the percentage of isolates carrying expressed TE copies. For instance, among DNA transposons, the MITE Undine had more than 50% of the copies expressed in the reference genome and more than 75% of the Undine copies were expressed among isolates. For the DTA hAT element, less than 30% of the copies were expressed in the genome as well as among isolates (Fig. 4D, Supplementary Fig. S4, Supplementary Table S6).

**Figure 4.**
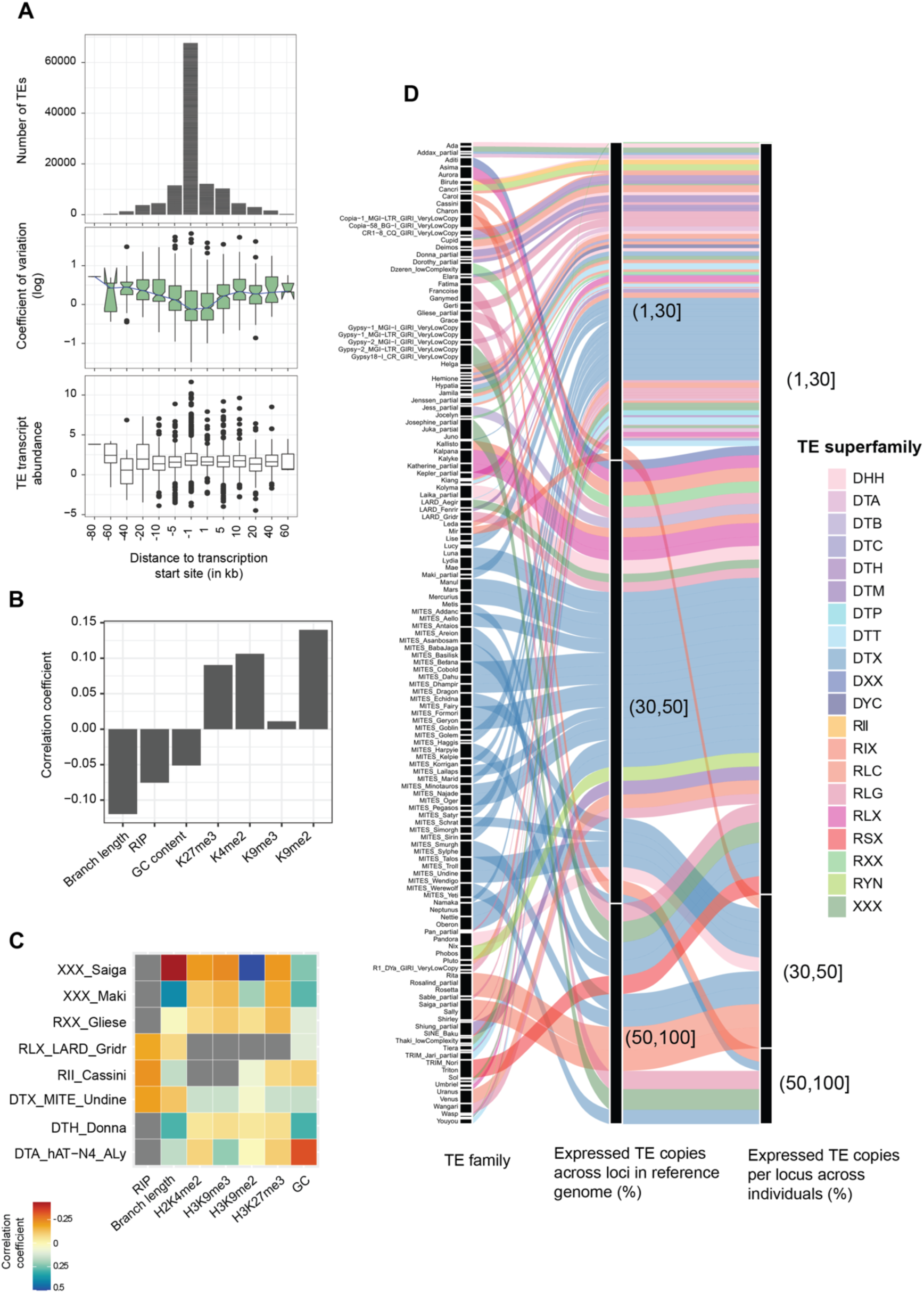
Transcriptional landscape of TEs across the genome. A) Number, transcriptional variation, and expression count of TE localized upstream and downstream of TSS of a gene B) Correlation of transcription of individual TE loci and corresponding genomic defense mechanism in the loci. C) Correlation of transcription of individual TE loci and corresponding genomic defense mechanism in the loci for TE families (TE families with > 25 expressed copies used for correlation) D) Percentage of expressed TE copies in the reference genome and percentage of expressed corresponding TE copies in the population.

### Enrichment of repeat-induced polymorphisms in regulatory regions of TEs

We used expression association mapping to systematically identify polymorphisms associated with variation in TE expression among isolates. We limited the search for associated SNPs to a 5 kb distance from the TE loci. We found 23 TE loci with at least one significantly associated SNP and the majority of the significant SNPs were located within 2 kb of the TE (Fig 5A-B). Next, we assessed whether there was an enrichment for RIP-like mutations among TE expression-associated polymorphisms. In *Z. tritici*, RIP introduces CèT transitions at CpA sites (Moller et al., 2021) and we observed an enrichment of CèT transitions in expression-associated polymorphisms at CpA sites (odds ratio of 2.70, CI 0.52 -13.98). In contrast, variants associated with gene expression showed no such bias (odds ratio of 1.03, CI 0.93-1.14; Fig 5C;). The effect size of RIP-like TE-associated regulatory polymorphisms was lower than the non-RIP-like regulatory polymorphisms (Fig 5D)

**Figure 5.**
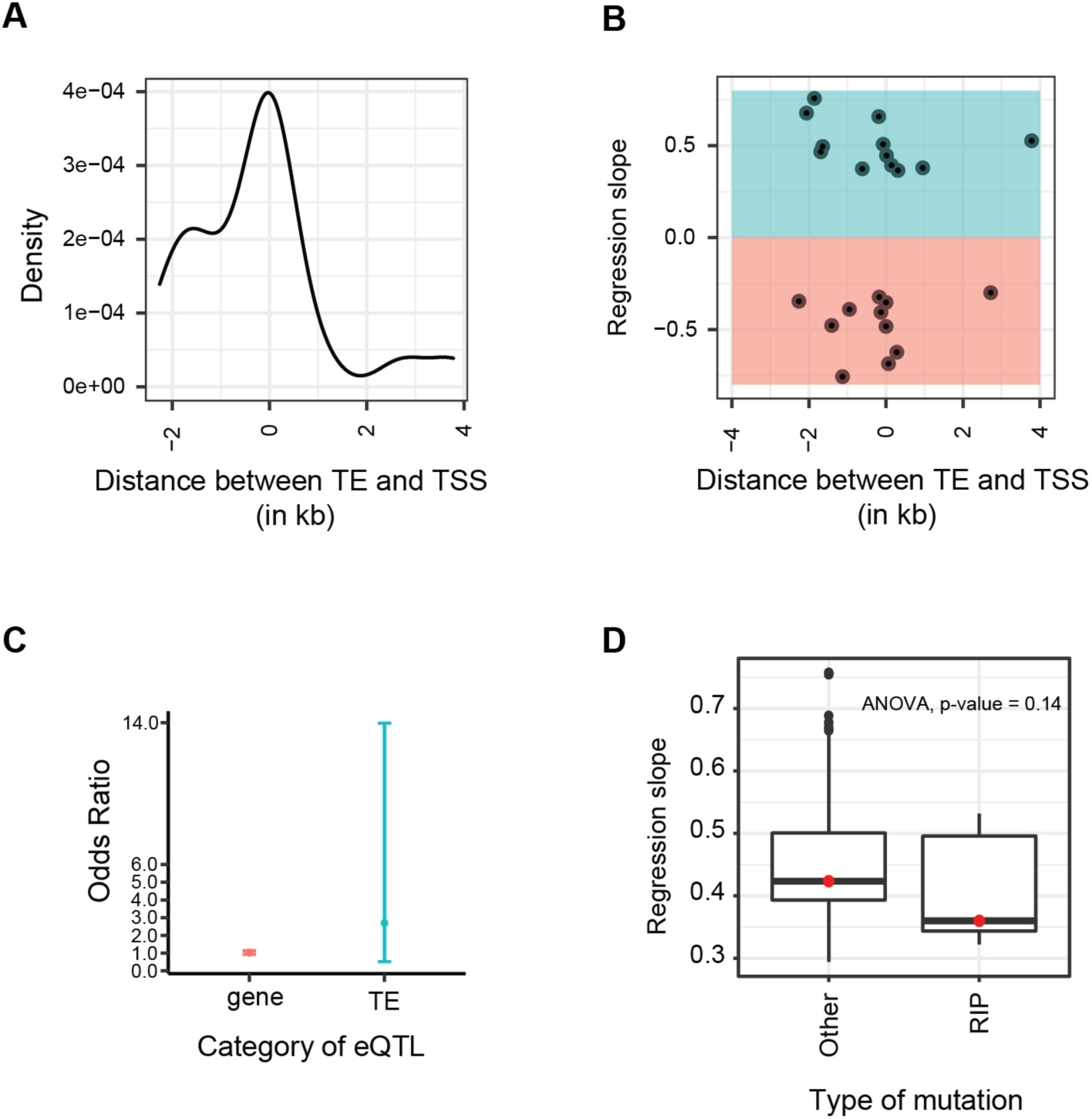
Regulatory variants associated with TE transcription. A) Distribution TE transcription-associated polymorphism within a 2 kb distance upstream and downstream of TE loci B) Effect size (*i.e.,* regression slope) of polymorphism associated with TE transcription within a 2 kb distance upstream and downstream of TE loci. C) Enrichment of RIP-like mutations in TE-transcription associated polymorphisms compared to gene-expression associated polymorphisms. D) Effect size of RIP-like and non-RIP mutations associated with TE transcription variation

### Phenotypic variation associated with recent TE insertions in a population

Insertions caused by TEs can have consequences for the expression of phenotypic traits. To investigate such effects within the field population, we first assessed patterns of linkage disequilibrium (LD) between TIPs and nearby SNPs. We found that most TIPs were on average in low LD with SNPs within a 600bp kb window compared to LD between SNPs in the same window (Fig 6A, Supplementary Table S7). However, the larger distance between SNPs and TIPs in the window compared to SNP-SNP distances explains most likely the low LD observed between SNPs and TIPs (Supplementary Fig S5). We also found that TIPs with a low-frequency TE insertion (<5%) had lower LD with neighboring SNPs compared to high-frequency TE insertions (Supplementary Fig S5). To identify associations of TIPs with phenotypic trait variation, we analyzed a series of trait datasets generated for the same population including pathogenicity trait measurements and variation in metabolite production (Supplementary Table S8). Wheat leaf lesion area produced by individual isolates showed a significant association (*p*-value*=*3.8*10-4) with a TIP caused by a 6 kb *Copia* element (RLC *Deimos*) located on chromosome 9 (Fig 6B). The associated TIP is within ∼5 kb of a gene encoding a cell wall-degrading enzyme (*i.e.,* glycosyl-hydrolase family 47) secreted during host infection. We found no significant difference in expression of the gene encoding the cell wall-degrading enzyme between isolates carrying the *Copia* element or not (*t*-test *p*-value = 0.44; Fig 6C).

**Figure 6.**
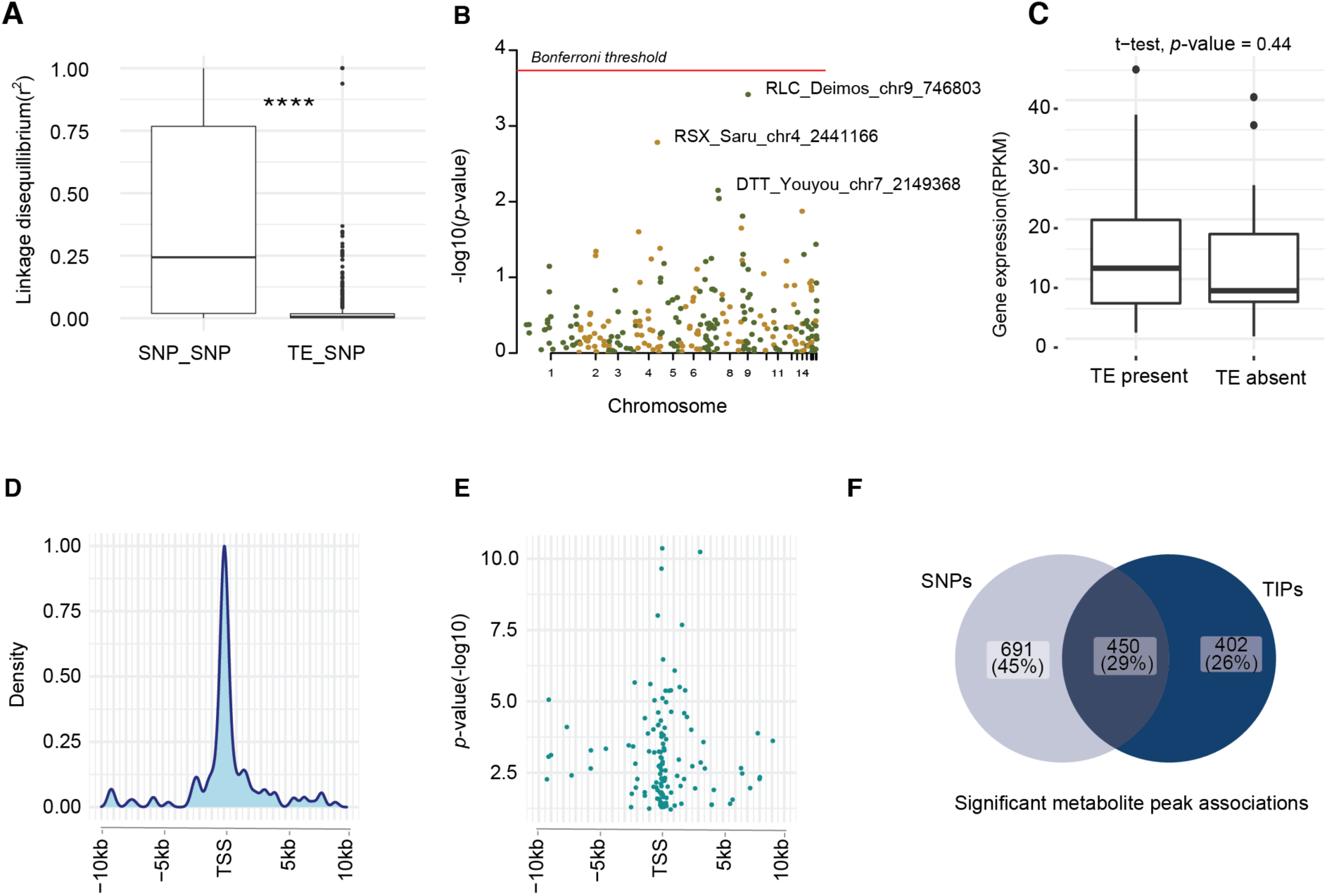
Phenotypic association of TIPs in the population. A) Linkage disequilibrium between TIPs and SNPs in the 5kb window and SNP-SNP LD in the 5kb window. B) Manhattan plot for TIP association with virulence (measured by percent leaf area covered by lesion) C) TIP and neighboring cell wall degrading enzyme gene expression difference. D) Distribution of TIP associated with metabolome variation around TSS of the gene. E) P value distribution of TIP associated with metabolome variation around TSS of gene F) A Venn diagram representing the overlap of significant association form TIP associated metabolome GWAS and SNP-associated metabolome GWAS

Given the large scope for TE insertions associated with gene transcription variation, we screened the population for additional phenotypic readouts. We analyzed a dataset of metabolite production profiles of each isolate assessed under culture condition (Singh et al., 2022). We found that 65% of all TIPs were significantly associated with variation in intensity in at least one metabolite (Supplementary Table S9). Out of the significantly associated TIPs, 13 were significantly associated with at least 20 different metabolites (Supplementary Fig 7). We found that TIPs located within ∼1 kb upstream of the transcription start site (TSS) of genes have a stronger association than TIPs further away from the TSS. Furthermore, significantly associated TIPs were enriched in the coding region and the TSS compared to TIPs located within 5 kb upstream and downstream of the TSS (Fig 6D,E). The enrichment of significant TIPs within coding regions and near TSS is consistent with the observations from a metabolome GWAS (Singh et al., 2022) performed on the same study population using genome-wide SNPs. Overall, 26% of all significant associations with individual metabolites using TIP-GWAS were not previously identified using SNPs only (Fig 6F).

**Figure 7.**
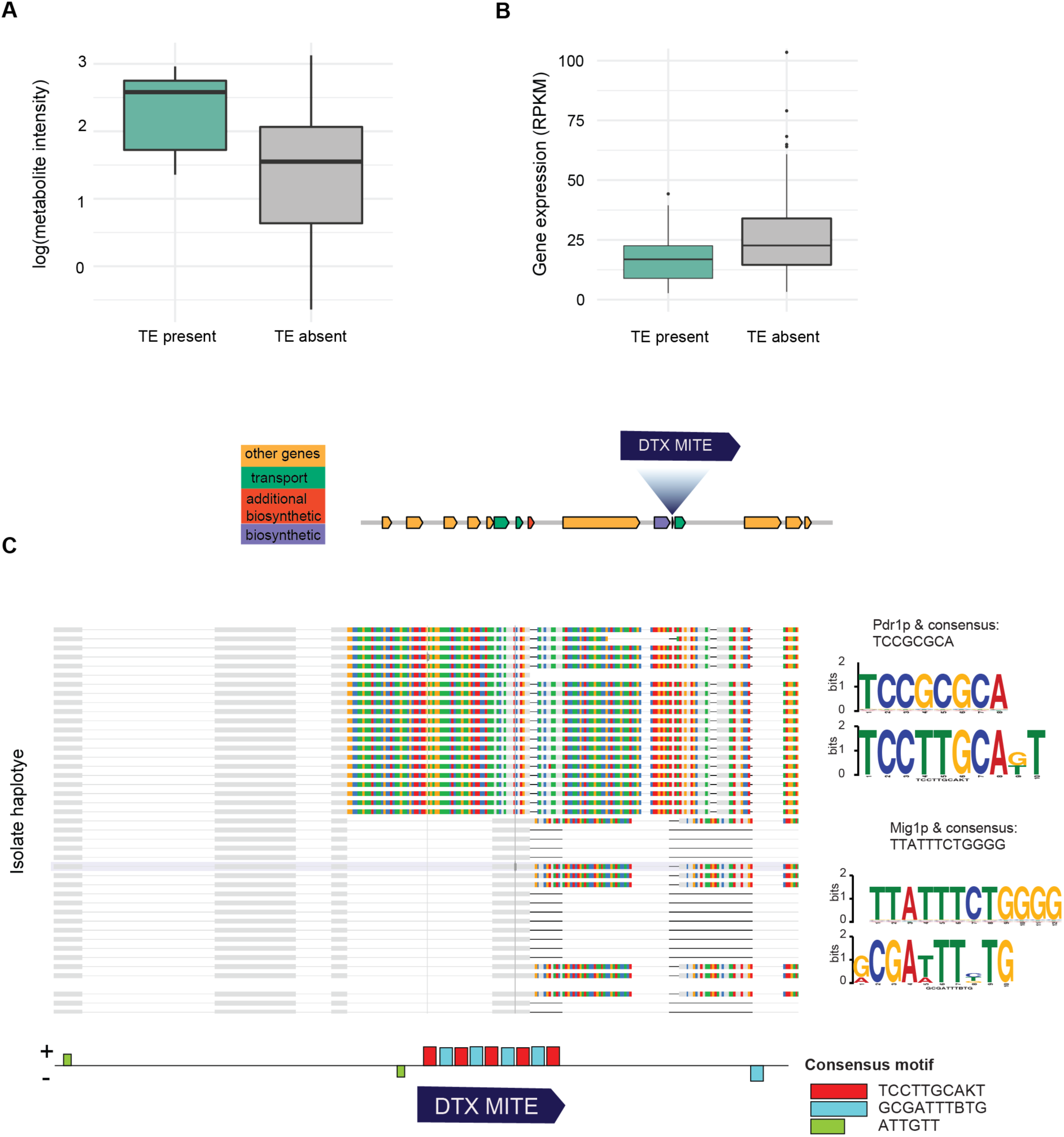
Genome context and enriched motifs near the MITE insertion in a secondary metabolite gene cluster. A) Association of metabolite production intensity with the MITE presence/absence variation. B) Association of the *NmrA*-like gene expression variation associated with the MITE presence/absence variation. C) Multiple sequence alignment of the intergenic region nearby the MITE insertion and motifs enriched near the insertion site.

Based on the metabolite-TIP GWAS, we identified a significant association for a DTX MITE insertion with the metabolite profile *m/z* intensity 245.1866. The associated MITE insertion is in the intergenic region of a polyketide synthase (PKS) secondary metabolite gene cluster associated with higher metabolite production (Fig 7A). The insertion occurred upstream of the gene encoding a NmrA-like family protein (Fig 7B). NmrA is a negative transcriptional regulator involved in nitrogen metabolite repression in pathogenic fungi (Li et al., 2021). We found that the MITE insertion was significantly associated with lower expression of the NmrA-like family protein (Fig 7B). To understand the molecular mechanism underpinning the downregulation of the *NmrA*-like family gene caused by the MITE insertion, we analyzed intergenic sequence variation including the TE insertion for motif enrichments. Two motifs in the TE sequence share similarities to binding sites of the transcription factor Pdr1p, a zinc cluster protein that can act as a master regulator involved in recruiting other zinc cluster proteins to pleiotropic drug response elements (PDEs) (MacPherson et al., 2006). PDEs fine-tune the regulation of multidrug resistance genes. The second motif shares sequence similarity with the binding site for Mig1p, a transcription factor involved in glucose repression in *Saccharomyces cerevisiae* (Fig 7C). The identified transcription factor binding motifs in the TEs suggest a regulatory role of MITE insertion and a beneficial effect on transcription factor binding.

## Discussion

We performed a comprehensive analysis of transcriptional variation generated by individual TEs in a pathogen population and consequences for phenotypic trait evolution. Recent TE insertions generated presence-absence variation among isolates (*i.e.,* TIPs) of a single field population with selection likely playing an important role in filtering new insertions. We identified considerable intraspecific variation in the transcription of individual TE copies emphasizing the importance of the genomic background for controlling TE activity. Finally, TE insertions likely influence the gene expression of neighboring genes and the expression of traits including metabolites, which can mediate organismal interactions.

We found approximately a fifth of all TE copies in the genome with transcriptional activity. This indicates significant potential for additional copies being created. Consistent with the expectation for new insertions being generated, a high number of TE insertions are not fixed within the population. Beyond the single field population, the species contains multiple, highly active TEs with diverse effects on trait expression, epigenetic variation, and control on genome size (Fouché et al., 2020; Oggenfuss & Croll, 2023; Torres et al., 2020). Highly active TEs likely create mutation load within species as proposed for *Arabidopsis thaliana* (Quadrana et al., 2019). With the large effective population size of *Z. tritici* (Hartmann et al., 2018), selection should act strongly against deleterious TE insertions. Furthermore, genomic defenses should be active to reduce the functional integrity of TE sequences. As expected from strong counter-selection, insertions are more likely to occur in non-coding regions than exon or intron sequences. We also found that the identity of TEs is an important factor explaining their distribution across gene elements with *e.g.,* some TEs nearly exclusively associated with intronic insertions. Some introns in *Z. tritici* were identified as non-autonomous TEs because unrelated genes across the genome harbor nearly identical intron sequences (Torriani et al., 2011; Wu et al., 2017). Overall, the species harbors dozens of polymorphisms for the presence-absence of entire intron sequences consistent with the recent activity of intronic sequences (Torriani et al., 2011). Retrotransposons were overrepresented in exonic sequences compare to DNA transposons. This may be explained at least partially by gene models including open reading frames of TEs or chimeric genes. Similar to patterns across the genome, the localization, and frequency of TIPs reflect selection acting against deleterious insertions. TE frequencies at TIPs tend to be low except the intronic TEs, which are most likely domesticated with no or only minor deleterious effects. Overall, the TE landscape of a single population strongly reflects insertion constraints and counter-selection acting against TE insertions.

Selection acting against deleterious TE insertions is complemented by specific genomic defense mechanisms that evolved to counteract TE activity in fungal genomes. These mechanisms include RIP mutations acting against duplicated sequences and epigenetic silencing reducing transcriptional activity of TE and accessibility of DNA (Gladyshev, 2017; Slotkin & Martienssen, 2007b; Young et al., 2020). We found mostly low to no transcription of TE copies affected by RIP suggesting that the defense mechanism is unable to effectively silence TEs as in *N. crassa* (Wang et al., 2020). The most abundant TE family (*i.e.*, the MITE Undine) showed the highest transcriptional variation for individual loci in the population. The same TE is also highly upregulated during plant infection and has undergone a massive recent expansion (Fouché et al., 2020; Oggenfuss & Croll, 2023). The weak effect of genomic defenses against this MITE is likely related to its short length, which renders RIP-based defenses ineffective (Fouché et al., 2020; Pereira et al., 2021). Active transcription of a TE can generate new TE insertions in the genome, as well as impact the structure and function of the genome (Feschotte, 2008; Wicker et al., 2018). The heterogeneous effects of RIP against different classes of TEs is compounded by the relaxation of RIP genomic defenses within the species following the expansion from the center of origin in the Middle East (Feurtey et al., 2023; Jovan Komluski, 2022; Moller et al., 2021).

The activity and spread in the genome were highly uneven for different TE families in the genome consistent with differences in recent activity, insertion preferences and evidence from studies across different kingdoms (Wells & Feschotte, 2020). The *Mutator* elements of maize typically integrate into gene-rich regions (Tan et al., 2011). In *S. cerevisiae*, *Copia/Ty1* integrates preferentially upstream of genes transcribed by the RNA polymerase III (Pol III) (Baller et al., 2012; Mularoni et al., 2012). *Ty1* is also enriched in heterochromatic region triggered by the recognition of heterochromatin during integration and then perpetuate the heterochromatic mark by triggering epigenetic modifications at new insertion loci (Gao et al., 2008). The *P* elements in *Drosophila* preferentially integrate at replication origins in the genome to support transposition (Spradling et al., 2011). We also find that TE insertions are underrepresented near highly expressed genes similar to the observation in *D. nasuta* (Wei et al., 2022b). Those observations may however be the result of a TE survivor bias given the typically slow evolutionary rates of highly expressed genes given the strong purifying selection (Drummond et al., 2005; Wei et al., 2022b). Under some conditions, the spread of heterochromatin from TE insertion sites can induce epigenetic silencing of neighboring genes (Young et al., 2020). Consistent with this possibility, we found that TE insertions are significantly associated with expression variation of neighboring genes. TE insertion tends to be associated with lower gene transcription levels consistent with previous analyses of gene expression analyses over the course of plant infections (Fouché et al., 2020). Based on environmental cues and stress induction, TEs show distinct de-repression patterns throughout an infection with correlated responses of genes in proximity (Fouché et al., 2019). Our findings expand our understanding of the coordinated expression of TEs and adjacent genes by showing that these patterns can be driven by polymorphic TE insertions.

Crucial for the adaptive evolution of species, TE insertion activity in genomes can drive phenotypic trait evolution through their impact on gene structure and regulation. We showed that variation in pathogenicity-related traits and metabolite production is most likely underpinned by recent TE insertion activity in functionally important regions of the genome. Our TIP-GWAS showed that two-thirds of all TIPs showed a significant association with at least one metabolite intensity profile. The specific biological roles of the different metabolites are poorly understood with few exceptions (Hassani et al., 2022; Singh et al., 2022b). However, individual metabolites can play major roles in species interactions with competitors or hosts (Derbyshire et al., 2018; Krishnan et al., 2018). Similarly, TE insertions in crop plants are associated with a wide range of agronomic traits and secondary metabolites such as in tomatoes (Domínguez et al., 2020). Polymorphic TE insertions also played a role in the selection of *Brassica rapa* morphotypes during domestication (Cai et al., 2022b). Metabolite production associated TIPs in *Z. tritici* showed an enrichment in coding regions and transcription start sites consistent with these regions having a higher potential for functional consequences. Given the large number of TE insertions associated across many genes for metabolite production, provides a vast potential for rapid evolution of the species from standing variation in single field populations. TE-driven adaptation within species and populations is likely to proceed at a more rapid pace given the potential for stronger phenotypic variation among genotypes at TIPs. Consistent with individual findings across plants, animals, and fungal species, active TEs in the genome can underpin most recent adaptive evolution of large effect size. The power of TEs to drive evolutionary change stems from their potential to affect the expression of multiple nearby genes, modulate their response to external stress but also to rearrange large sequence segments including the deletion and duplication of gene sets. Recent discoveries of large TEs with the ability to conjugate dozens of genes and hundreds of kilobases of sequences (Gluck-Thaler et al., 2022) exemplifies the driving force selfish elements play underpinning rapid evolution of species.

## Acknowledgments

RNA-seq data generated for this manuscript was obtained in collaboration with the Genetic Diversity Centre (GDC), ETH Zurich, and the iGE3 platform of the University of Geneva.

## Data availability

All RNA-seq datasets are available from the NCBI Sequence Read Archive BioProject PRJNA596434. Metabolite profile variation is available from the Supplementary Information of Singh et al. (2022). Supplementary Tables provide all TE insertion, expression and association mapping data.

### Author contributions

LNA and DC conceived the study. LNA generated the RNA-seq data. UO generated the TE polymorphism data. LNA performed data analyses. LNA wrote the manuscript with DC.

