## Supplementary Figures for "Population-level transposable element expression dynamics influence trait evolution in a fungal crop pathogen"

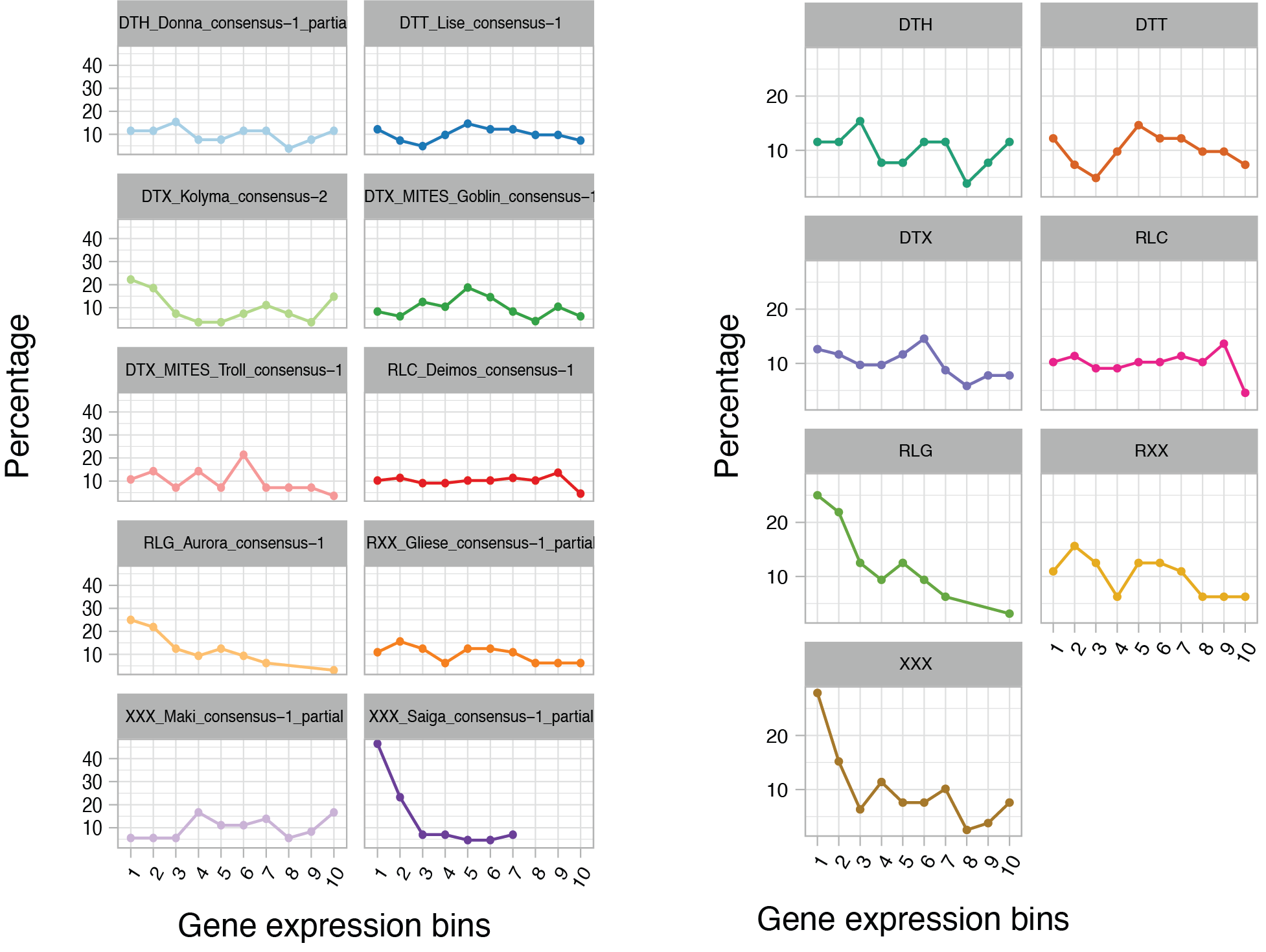

**Supplementary Figure S1**: Distribution of TE copies of individual families and grouped by TE superfamily across genes ranked by gene expression. Gene expression bins refer to genes ordered by increasing order of gene expression (RPKM).

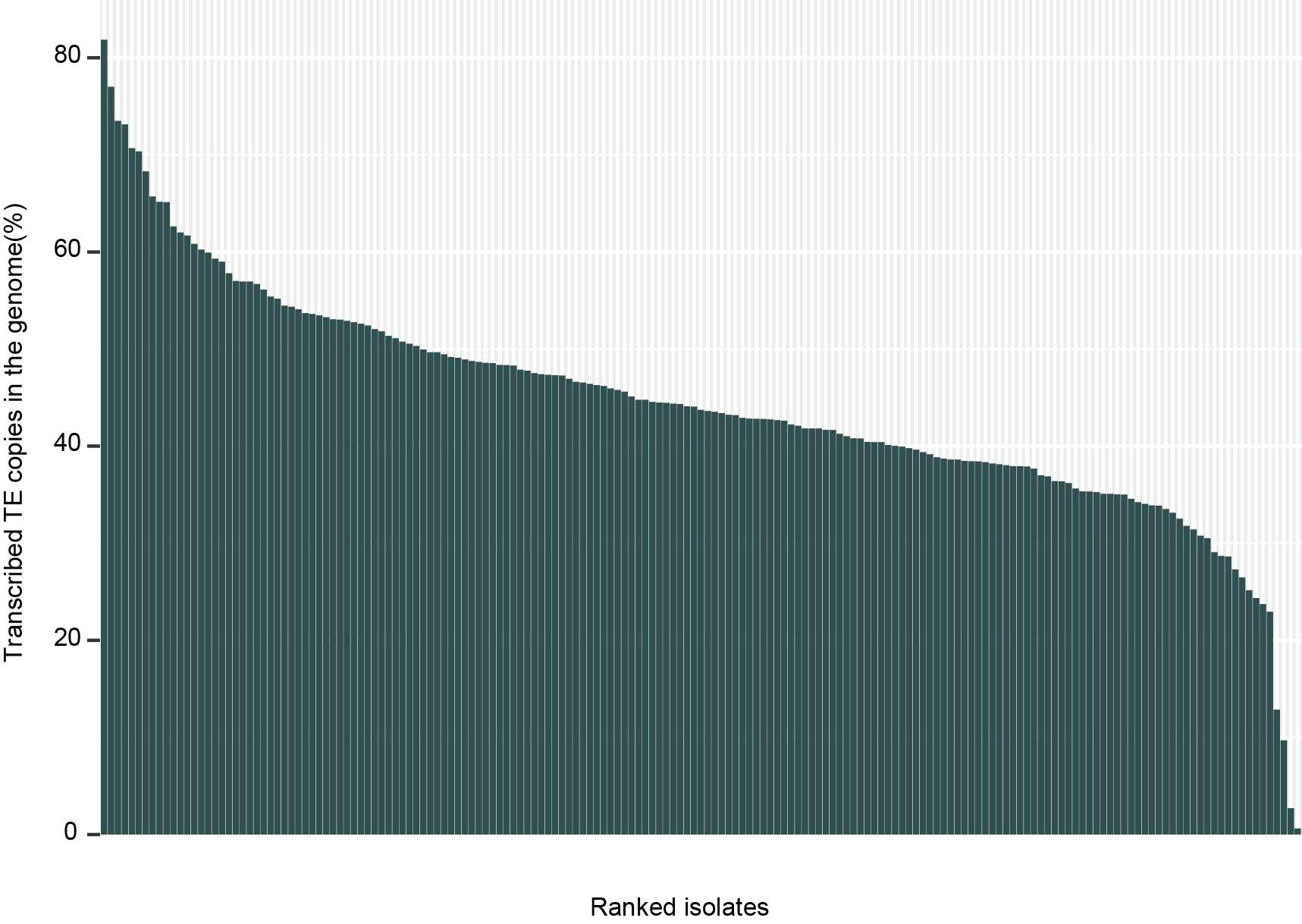

**Supplementary Figure S2**: Percent of transcribed TE copies in the genome of isolates in the population.

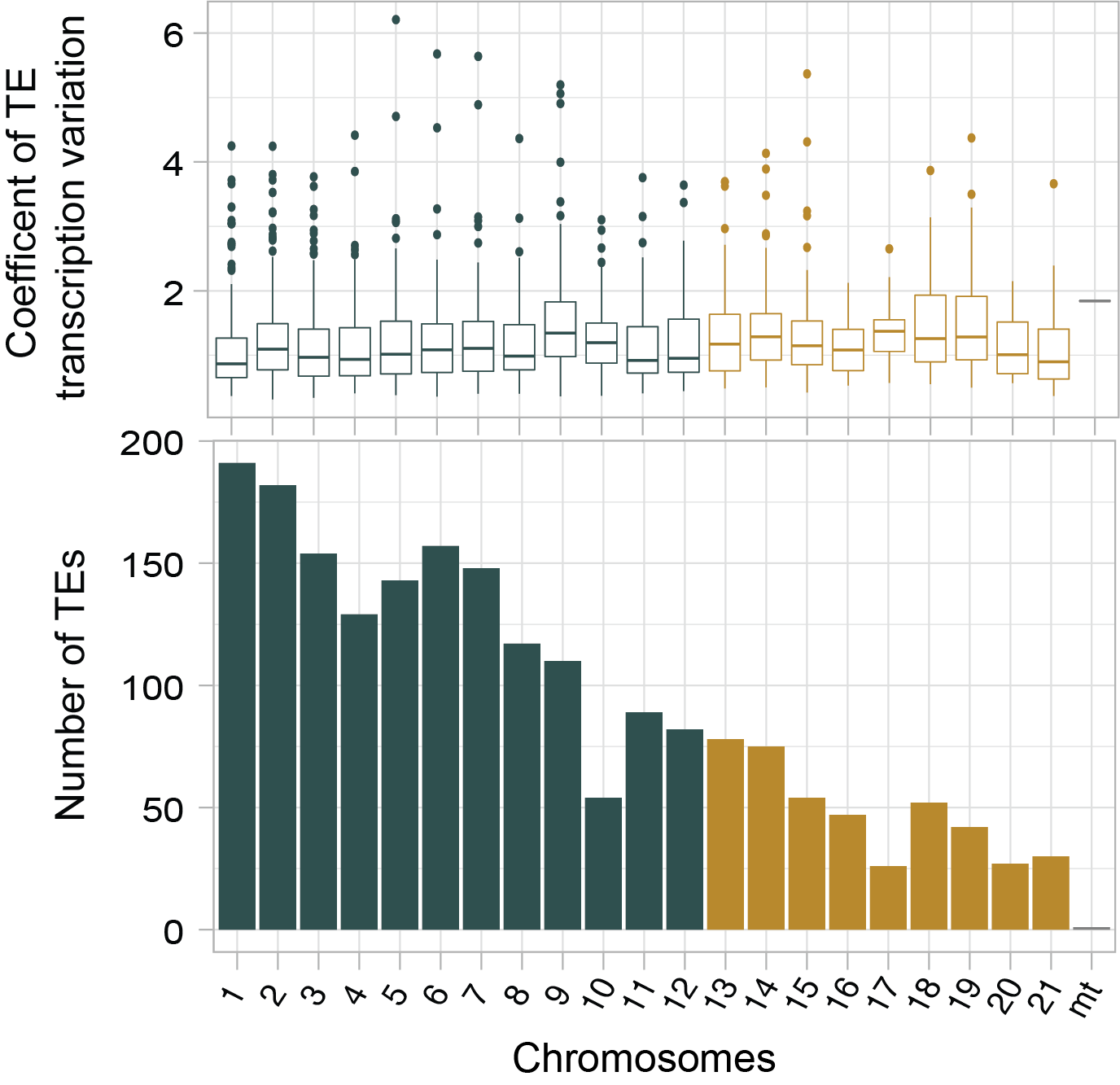

**Supplementary Figure S3**: Population-level transcriptional variation of individual TE copies binned by chromosome. Distribution of TE copies across chromosomes.

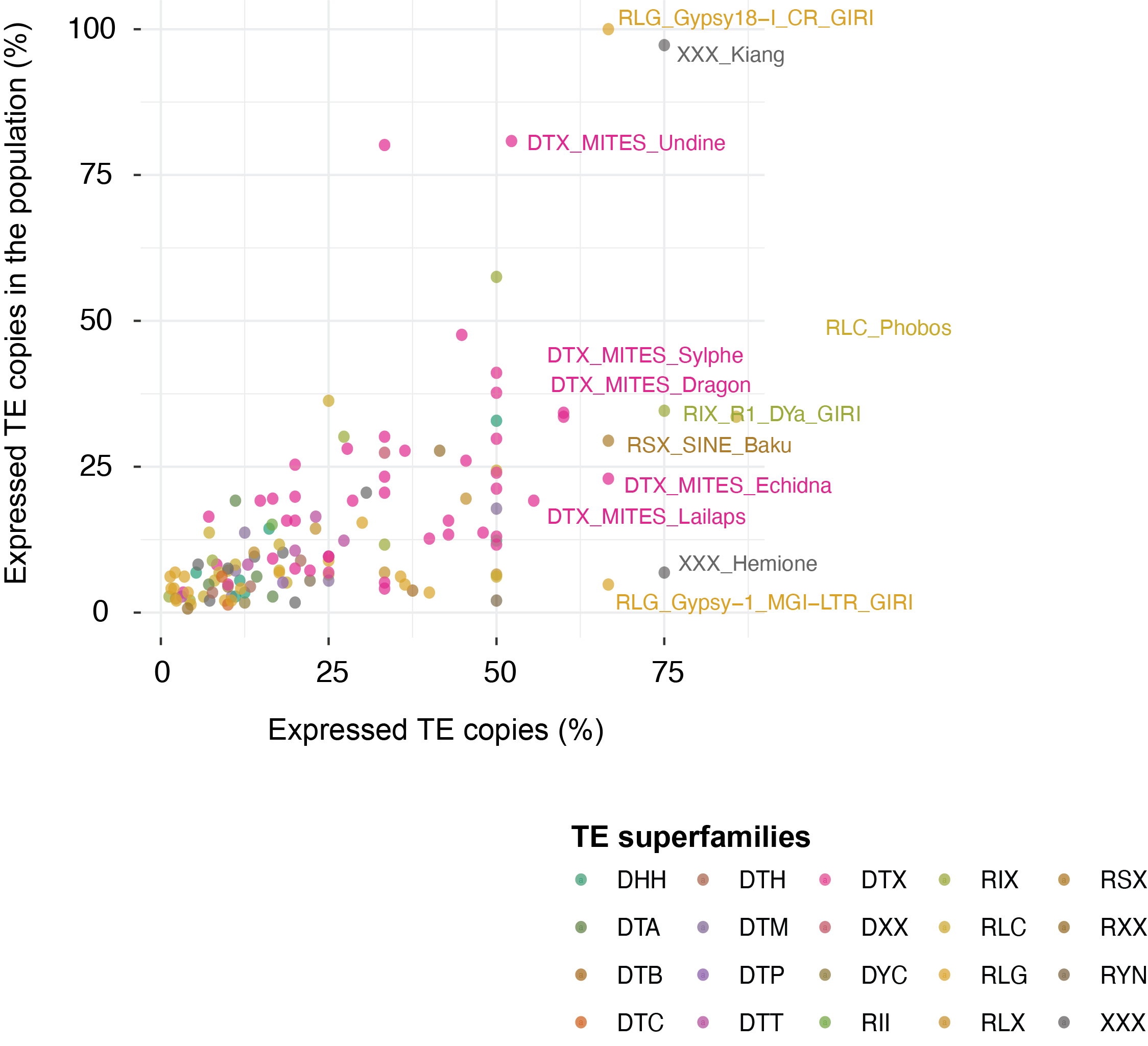

**Supplementary Figure S4**: Percent of expressed TE copies in the reference genome isolate IPO323 and percent of TE copies expressed across the population. Each dot represents a TE family and the percentages refer to different copies of the same family. Colors indicate superfamily groups.

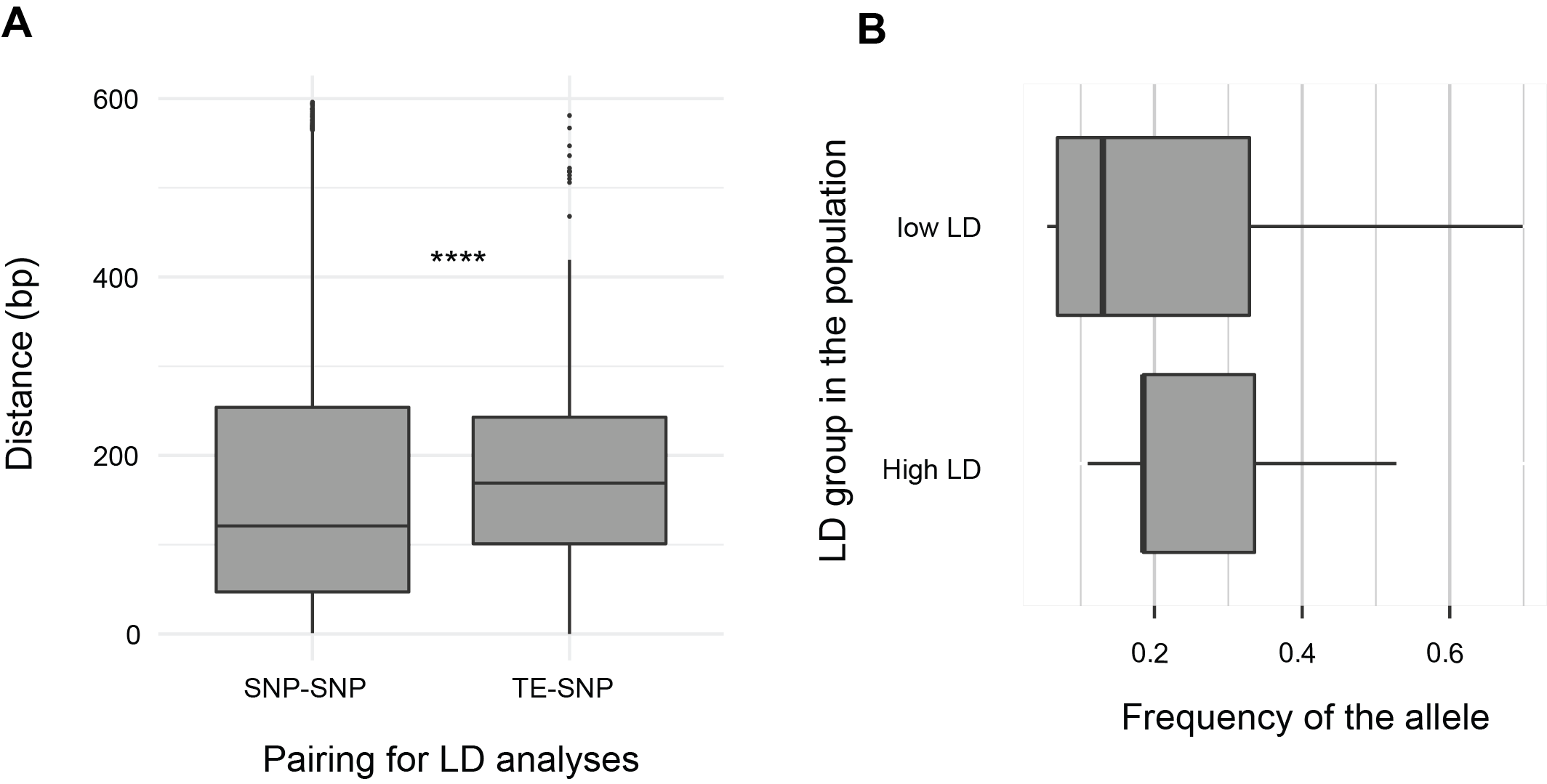

**Supplementary Figure S5**: (A) Distances between pairs of SNPs, and pairs of TIPs with SNPs in the 5 kb upstream and down steam from a focal TIP position. (B) Frequency of the TIPs in the population and linkage disequilibrium (LD) with the neighboring SNPs in the 5kb windows up- and downstream of each focal TIP.

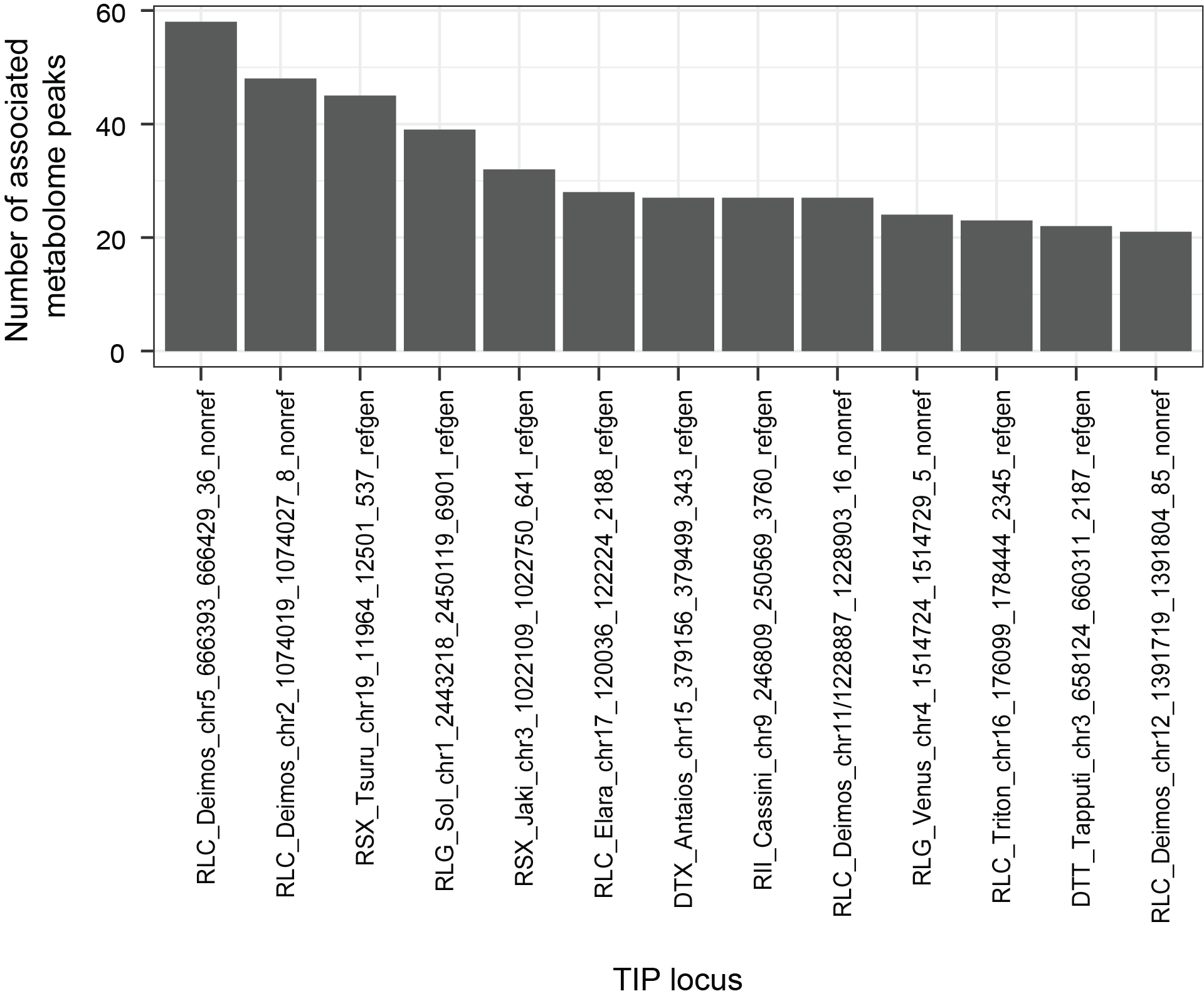

**Supplementary Figure S6**: TIPs significantly associated with variation in ≥20 metabolite peak profiles.

**Supplementary Tables**

[see separate file]

**Supplementary Table S1:** SRA accession list of RNAseq reads.

**Supplementary Table S2:** Genomic localization of TEs in gene elements and 10 kb windows upstream and downstream of the transcription start site (TSS) in the reference genome IPO323.

**Supplementary Table S3:** Genome-wide TE insertion polymorphism (TIPs) in the pathogen population. 0 represents TE absence and 1 represents TE presence.

**Supplementary Table S4:** Gene expression (log-transformed RPKM) values across the population.

**Supplementary Table S5:** Locus-specific transcript abundance at individual TE loci (FPKM) across individuals.

**Supplementary Table S6:** Percent of expressed copies within each TE family in the reference genome IPO323 and percent expressed TE copies in each TE family across the population.

**Supplementary Table S7:** Linkage disequilibrium of TIP in the genome and neighboring SNPs within the 600bp distance from the TE loci.

**Supplementary Table S8:** Genome-wide association mapping of the virulence-associated trait (PLACP: percent leaf area covered by pycnidia) and TE insertion polymorphisms in the genome.

**Supplementary Table S9:** TIPs in the genome significantly associated with metabolite peak intensity variation in the pathogen population (filtered by Bonferroni threshold).
